# Epigenetic clock and methylation studies in dogs

**DOI:** 10.1101/2021.03.30.437604

**Authors:** Steve Horvath, Ake T. Lu, Amin Haghani, Joseph A. Zoller, Robert T. Brooke, Ken Raj, Jocelyn Plassais, Andrew N. Hogan, Elaine A. Ostrander

## Abstract

DNA methylation profiles have been used to develop biomarkers of aging known as epigenetic clocks, which predict chronological age with remarkable accuracy and show promise for inferring health status as an indicator of biological age. Epigenetic clocks were first built to monitor human aging but the principles underpinning them appear to be evolutionarily conserved, as they have now been successfully developed for over 120 mammalian species. Here we describe reliable and highly accurate epigenetic clocks shown to apply to 51 domestic dog breeds. The methylation profiles were generated using a custom array with DNA sequences that are conserved across all mammalian species (HorvathMammalMethylChip40). Canine epigenetic clocks were constructed to estimate age and also predict mortality risk by estimating average time-to-death. We also present two highly accurate human-dog dual species epigenetic clocks (R=0.97), which may facilitate the ready translation from canine to human use (or vice versa) of anti-aging treatments being developed for longevity and preventive medicine. Finally, epigenome-wide association studies herein reveal individual methylation sites that may underlie the inverse relationship between breed weight and lifespan. Overall, we describe robust biomarkers to measure aging and potentially health status in canines.

## INTRODUCTION

While rodent models have led to many advancements in human health and biology, they also have well known shortcomings. Many new anti-aging interventions are currently in development worldwide and, ideally, follow-on testing will occur in species that are evolutionarily closer to humans, more similar in size, have high genetic diversity, and share the same environment as humans. Domestic dogs fulfill most of these criteria, offering a unique opportunity to evaluate the effectiveness of emerging anti-aging interventions^1–4^ There is also a significant need for health and medical monitoring tools for this popular pet, as there are more than 75 million dogs in the U.S. alone.

There are over 340 recognized dog breeds worldwide, each a closed breeding population under strong selection for morphologic and behavioral traits. As a result, dogs share extensive phenotypic and genetic homogeneity within breeds, and increased heterogeneity between breeds^5^. Small breeds live considerably longer than large breeds, offering the rare chance to understand the relationship between size and lifespan within a single mammalian species. Dogs also share a similar level of medical scrutiny as humans, experiencing the same hallmarks of aging and development including infancy, puberty, adulthood, old age, etc., but do so in about 20% of the time as humans^4,6^ As a result, dogs represent an ideal system for studies of comparative aging, where within breed studies can be conducted in a background of limited diversity, while between breed studies recapitulate the levels of diversity observed in humans.

Our previous presentation of an epigenetic clock for dogs and wolves^7^ represented one of the first non-human mammals for which an epigenetic clock was developed, and suggests that an evolutionarily conserved mechanism underlies DNA methylation across all mammals. This work shows that the age dependence of DNA methylation is conserved at syntenic sites in the genomes of multiple mammalian species including humans. However, generalizability of results was limited by low sample size (N<150) as well as technical limitations associated with the measurement platform (reduced representation bisulfite sequencing). Further, our initial study utilized only a few canine breeds which prevented testing the relationship between epigenetic aging and breed lifespan. Here we developed a new canine epigenetic clock based on 51 recognized dog breeds,^8^ and using a custom array that profiles cytosines whose flanking DNA sequences are conserved across mammalian species^9^

In this study, we aim to develop robust epigenetic clocks based on dog breeds that apply to both humans and dogs. We test, specifically, if short-lived breeds exhibit faster epigenetic aging rates than large breeds. We develop mortality risk estimators that model average time to death. Finally, we investigate the relationship between breed size and lifespan, and characterize CpGs with methylation changes that correlate with age, breed lifespan, and breed average adult weight, testing for changes that affect lifespan and weight separately and together.

## RESULTS

### DNA methylation data

All DNA methylation data were generated using the mammalian methylation array (HorvathMammalMethylChip40) that measures cytosine methylation levels in highly conserved regions across mammals^9^ We analyzed methylation profiles from 565 blood samples derived from 51 dog breeds *(Canis lupus familiaris*). Primary characteristics (sex, age, average life expectancy) for the breeds utilized are presented in **Supplementary Table 1**. The 51 breeds ranged from Bernese mountain dog, with the shortest expected lifespan of eight years (average adult breed weight=41 kg) to Chihuahuas with the highest expected lifespan at 20 years (average adult breed weight=1.8 kg). Expected lifespan estimates were based on the upper limit of lifespan estimated by the American Kennel Club and other registering bodies using sex averaged measures^10^. Unsupervised hierarchical clustering demonstrates that methylation profiles grouped by sex above breed (**Supplementary Figure S1**), i.e., most females were in one branch while most males grouped in a second branch. We did not find any evidence of grouping by breed. An attempt to build a methylation-based classifier of dog breed was unsuccessful (unreported data).

Using the same 565 samples, we then trained DNA methylation-based age estimators (epigenetic clocks). To generate dual-species (human-dog) clocks, we added 1,207 human DNA methylation profiles derived from the same custom array, which measures methylation levels at 37,492 cytosines with flanking DNA sequences that are conserved across mammalian species. From these, we produced two different epigenetic clocks that apply to both humans and dogs (**Methods**).

### Epigenetic clocks for dogs

The dog-only clocks were developed using blood DNA, while the human DNA that was used to generate the human-dog clocks were either from blood or multiple human tissues. The distinction between the two human-dog clocks lies in measurement parameters. One estimates *chronological* age (in units of years), while the other estimates *relative* age, which is the ratio of age of an individual to the maximum recorded lifespan of the species; with values between 0 and 1. This ratio allows alignment and biologically meaningful comparison between species with very different lifespans (dog and human), which is not afforded by the simple measurement of chronological age.

The cross-validation study reports unbiased estimates of the age correlation R, defined as Pearson correlation between the age estimate (DNAm age) and chronological age, as well as the median absolute error. Cross-validation estimates of age correlation for the three dog clocks are 0.97 (**Figure 1a, b, f, h**). Different cross validation schemes show that both the pure dog clock and the human-dog clock for chronological age exhibit a median error of less than 0.57 years (seven months) when using blood samples from dogs (**Figure 1a, b, f, i**).

**Figure 1:**
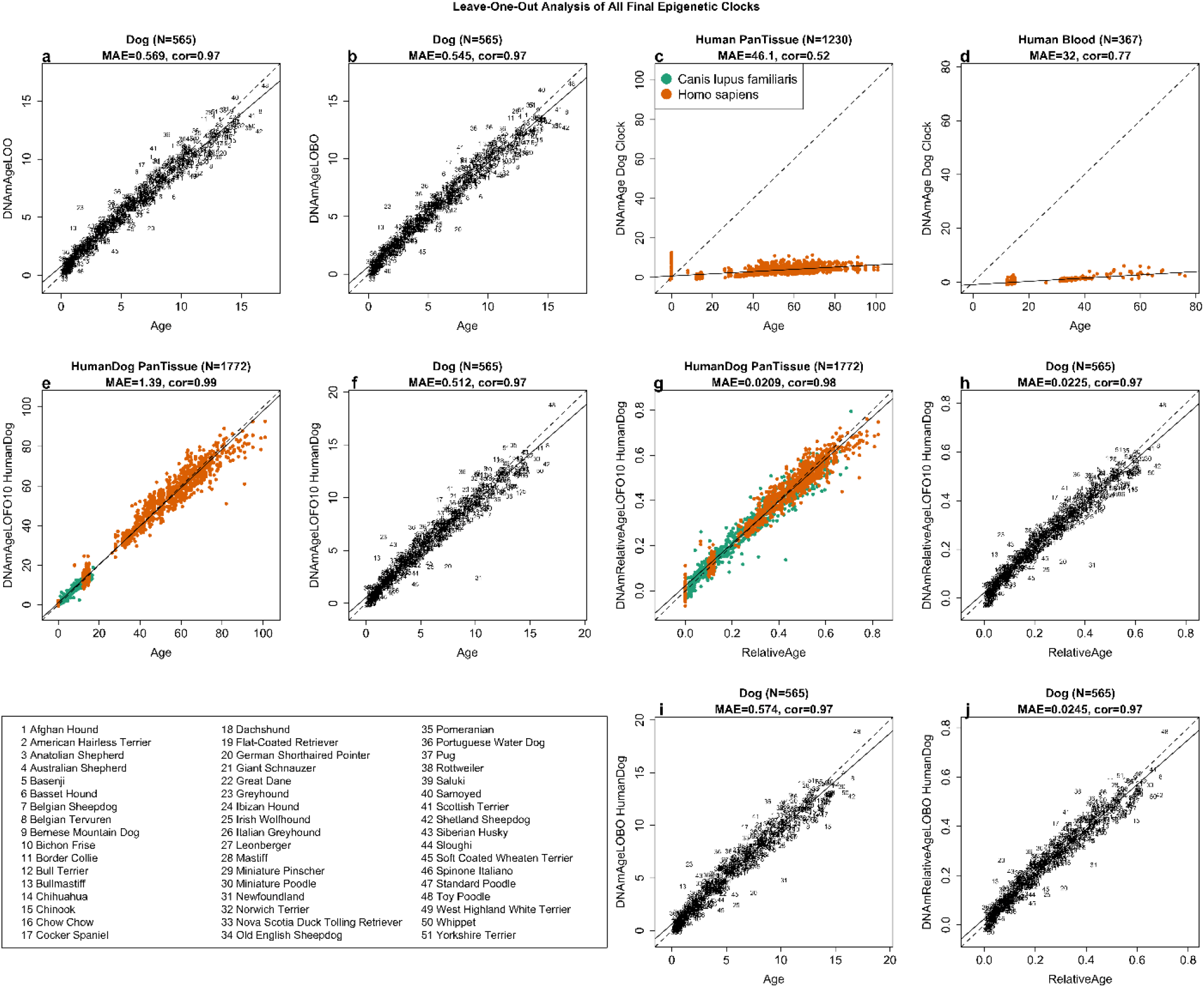
Cross-validation study of epigenetic clocks for dogs. To arrive at unbiased estimates of the dog epigenetic clock we carried out two types of validation studies: a) leave one out (LOO) cross validation and, b) leave-one-breed-out cross validation (LOBO). c, d) performance of the dog clock in c) different human tissues and d) human blood tissue. e,f,g,h) Species balanced cross validation (LOFO10Balance) analysis of human dog clocks for e,f) chronological age and g,h) relative age. Human dog clock for chronological age applied to e) both species and f) dogs only. Human dog clock for relative age applied to g) both species, h) dogs only. i,j) LOBO cross validation of the human dog clock for i) chronological age and j) relative age in blood samples from dogs. Species balanced cross validation (LOFO10Balance) was implemented in the following steps. First, we partitioned both the combined human/dog dataset into 10 evenly sized folds, where each fold has the same proportion and human and dog samples (referred to as “balanced” folds). We then iterate through each fold, training on the other nine folds, and applied the model to the target fold. Each panel reports the sample size, correlation coefficient, median absolute error (MAE).

By definition, the pure dog clock is not expected to apply to human tissues. However, we observe a remarkably high age correlation in DNA from human blood samples (r=0.77), albeit with a large median error of 32 years (**Figure 1d**). The age correlation of the dog clock across a variety of human tissues is lower (r=0.52, **Figure 1c**). By contrast, the two human dog clocks are highly accurate in both species (**Figure 1e-j**). The human-dog clock for *chronological* age led to a high age correlation of R=0.99 when both species are analyzed together (**Figure 1e**) and remained so when the analysis was restricted to dog samples alone (R=0.97, **Figure 1f**). Similarly, the human-dog clock for *relative age* exhibits a high correlation regardless of whether the analysis is done with samples from both species (R=0.98, **Figure 1g**) or only from dogs (R=0.97, **Figure 1h**). The impressive accuracy of the human-dog clocks could also be corroborated with an alternative cross validation scheme, i.e., a “leave one dog breed out” (LOBO) cross validation scheme (**Figure 1i, j**), which estimates the clock accuracy in dog breeds not used in the training set.

### Epigenetic clock to predict average time-to-death

Although epigenetic age clocks can be indirectly employed to predict risk or mortality, their performance may be sub-optimal, as they were developed for the clear purpose of estimating age^11,12^. The DNA methylation data that we generate, however, can be used to develop an epigenetic predictor of average time to death (“DNAmAverageTimeToDeath”), using a penalized regression model (**Methods**). To evaluate predictive accuracy, we used LOBO crossvalidation that divided the dataset into 51 breed groups. At each round, LOBO cross-validation trained each model on all but one breed, which was left out and used for validation at each iteration. LOBO analysis revealed a high correlation (r=0.89. **Figure 2a**) between estimated and actual average time to death with a median absolute error (MAE) of 1.3 years. As the successful performance of the LOBO analysis was carried out with 51 distinct breeds, we hypothesize that this model can be extrapolated to breeds not included in our dataset.

**Figure 2.**
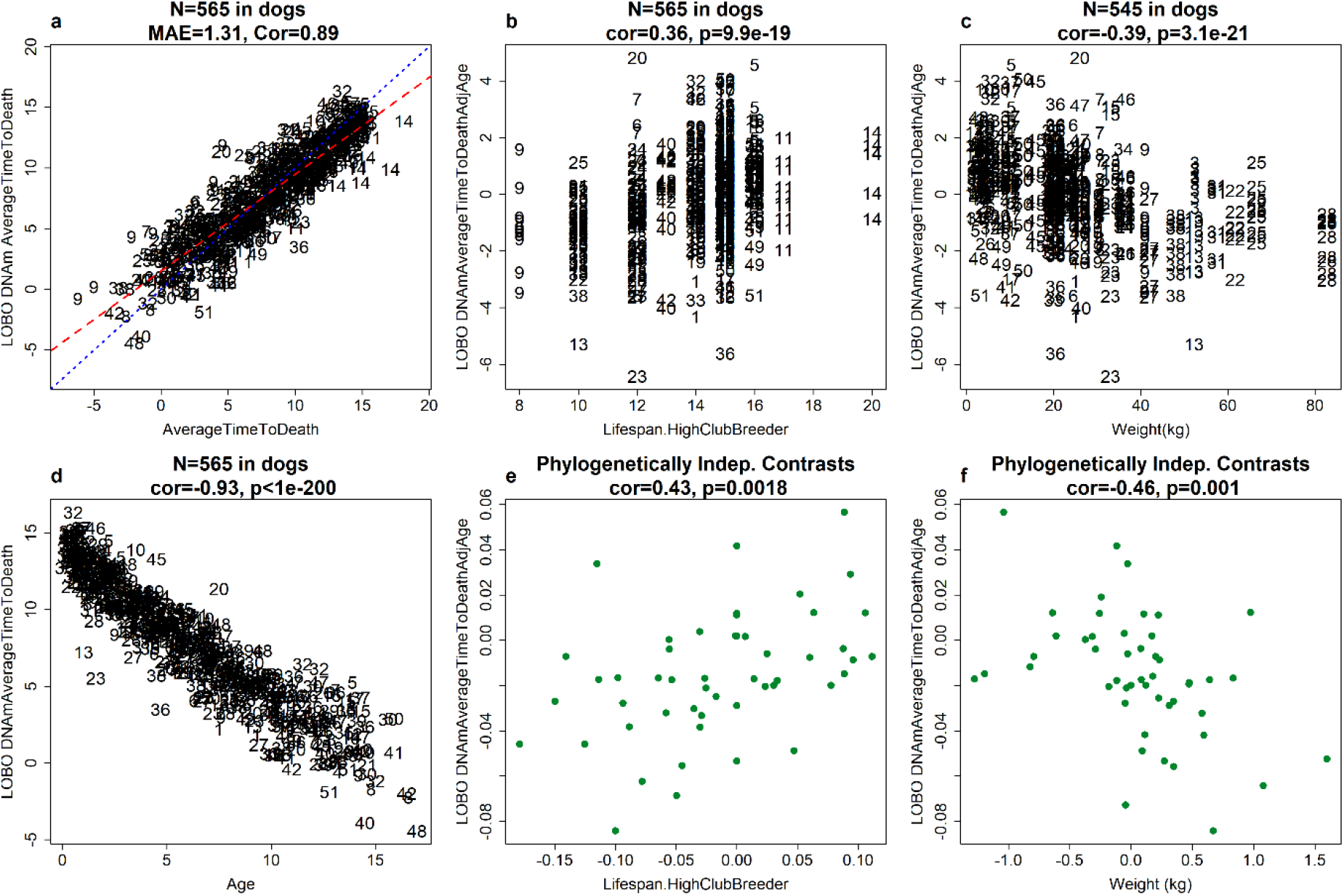
Epigenetic clocks for prediction average time to death. a) Leave-one-breed-out (LOBO) estimates of DNA methylation (DNAm) average time to death (y-axis, in units of years) versus average time to death (x-axis in units of years). For each dog, the average time to death was defined as difference between the upper limit of the respective breed lifespan (Lifespan.HighClubBreeder) and chronological age. b) LOBO DNAm average time to death adjusted for age (y-axis) versus lifespan (x-axis, in units of year). c) LOBO DNAm average time to death adjusted for age (y-axis) versus adult weight (x-axis). d) LOBO DNAm average time to death(y-axis) versus chronological age (x-axis). The association between LOBO DNAm average time to death adjusted for age and the lifespan remains significant (p= 4.4×10^-3^) even after adjusting for average adult weight in a multivariate regression model. e) Phylogenetically Independent (Indep.) contrast (PIC) generated LOBO DNAm average time to death adjusted for age (y-axis, at breed level) versus PIC generated lifespan (x-axis, at breed level). f) PIC generated LOBO DNAm average time to death adjusted for age (y-axis, at breed level) versus PIC generated adult weight (x-axis, at breed level). At each panel, we report Pearson correlation estimate. Each individual dog was marked in breed index as listed in the legend of Figure 1.

We observed that age-adjusted DNAmAverageTimeToDeath was correlated in the expected direction with lifespan (r=0.36 and P=9.9×10^-19^ **Figure 2b**) and average adult breed weight (r= −0.4 and P=3.1×10^-21^, **Figure 2c**). To account for correlated evolution of traits (lifespan, adult weight, etc.) among breeds, we applied phylogenetically independence contrasts (PICs) analysis to our study traits at the breed level. Age-adjusted DNAmAverageTimeToDeath retained similar degree of correlation with lifespan (r=0.43 and P=1.8×10^-3^, **Figure 2e**) and weight (r=-0.46 and P=1.0×10^-3^, **Figure 2f**), respectively. As expected from its construction, DNAmAverageTimeToDeath has a strong negative correlation with chronological age (r=0.93, **Figure 2d**). This is consistent with the fact that younger dogs are many more years away from reaching the upper limit of their lifespan for their respective breed. A multivariate regression model shows that the predictive accuracy of DNAmAverageTimeToDeath is retained even after adjusting for chronological age, gender and the average adult weight of the breed (P=1.56×10^-3^, **Table 1**).

**Table 1:** Linear regression analysis for dog breed lifespan

| **Model 1: dependent variable: lifespan** | **$\beta$** | **SE** | **T**<br>**statistic** | **P** |
| --- | --- | --- | --- | --- |
| Intercept | 15.882 | 0.136 | 116.385 | <2.0E-16 |
| Age-adjusted DNAmAverageTimeToDeath | 0.139 | 0.044 | 3.179 | 1.56E-03 |
| Female | -0.029 | 0.126 | -0.233 | 8.00E-01 |
| Weight (in units of kg) | -0.077 | 0.004 | -19.449 | 2.80E-64 |
| **Model 2: dependent variable: PICs lifespan** |  |  |  |  |
| PICs Age-adjusted DNAmAverageTimeToDeath | 0.566 | 0.312 | 1.816 | 0.076 |
| PICs weight | -0.055 | 0.018 | -3.140 | 2.95E-03 |

Note: Top panel presents the results from linear regression model analysis of dog breed lifespan (dependent variable) on age-adjusted DNAm average time to death(in units of years), gender and breed weight. The lower panel presents the regression analysis based on phylogenetically independent contrasts (PICs) generated traits. PICs traits are residuals that adjust the correlation of evolution traits. We set up the intercept term at zero, accordingly. We find suggestive evidence ( $P=0.08$ , Table 1) that DNAmAverageTimeToDeath continues to be associated with average time to death after adjusting for phylogenetic relationships between the dog breeds and age. The columns indicate the covariate name, regression coefficient, standard error, Student T-statistic, and two-sided Wald test p-value.

### EWAS of age and breed lifespan

In total, 31,911 probes of the mammalian methylation array are aligned to loci that are proximal to 5,021 genes in the recently released long read reference assembly of the Great Dane (CanFam_GreatDane.UMICH_Zoey_3.1.100)^13^. The array has high inter-species conservation; thus, it can be extrapolated to other breeds and other mammalian species^9^. As expected, age had a strong effect on DNAm levels in dogs and showed a similar aging biology to many previously analyzed mammalian species^14^. At p<10^-8^, the methylation level of a subset of 12,618 (40% of total) CpGs were related to age (**Figure 3a**). Top age-related CpGs in dog were located in the intron of *SLC12A5* (correlation test Z statistic, z = 42), the promoter of *MARCH4* (z = 37), and an intron of *LHX2* (z = 32). In general, CpGs that gained methylation with age were located near polycomb repressor complex 2 targets and enriched with genes that play a role in development (**Supplementary Figure S3**). Promoters largely gained methylation with age (**Figure 3b**).

**Figure 3.**
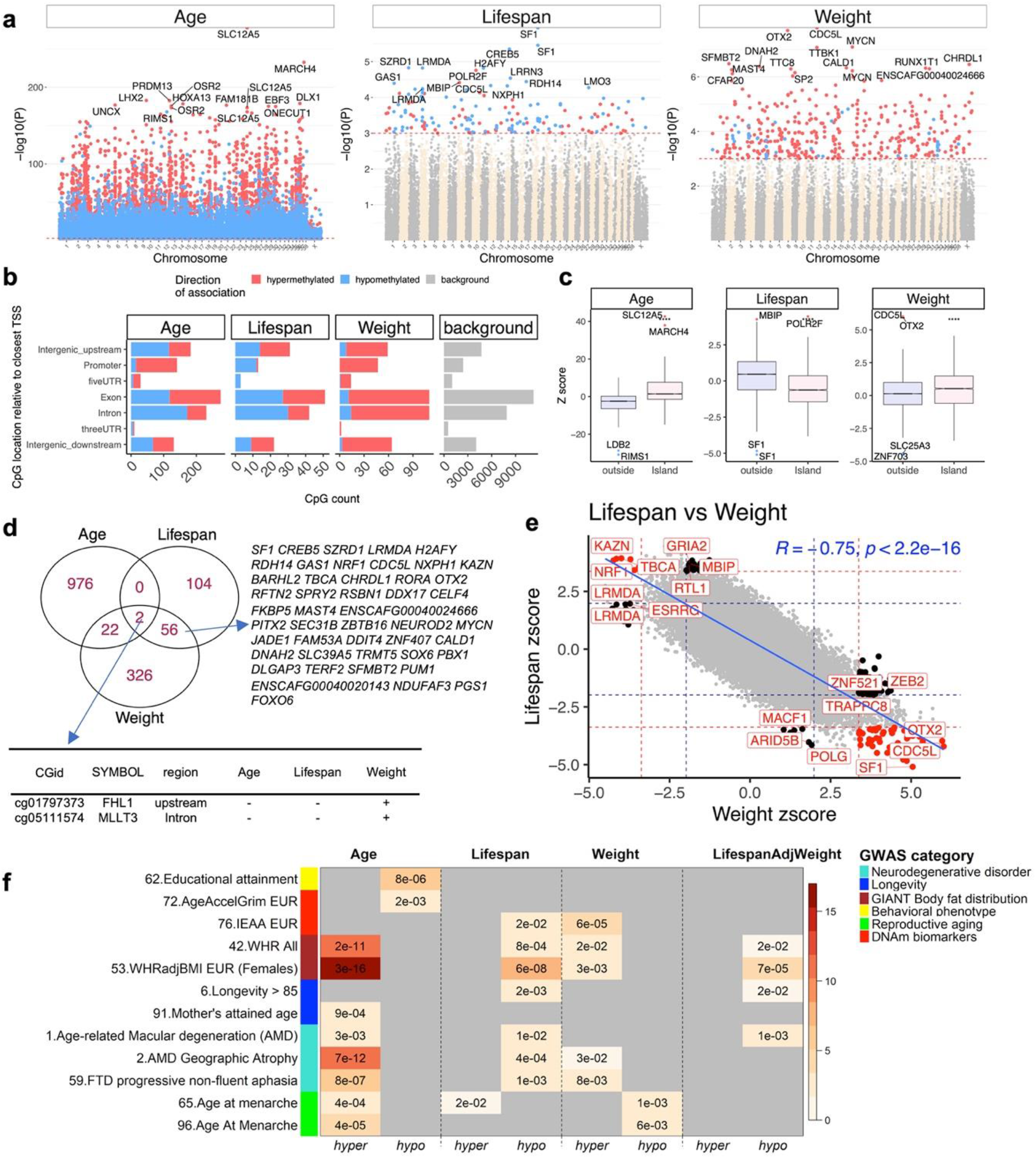
Epigenome wide association analysis of chronological age, average breed lifespan, and average breed weight of dog blood. a) Manhattan plots of the EWAS results. Coordinates are estimated based on the alignment of the mammalian array probes to the CanFam_GreatDane.UMICH_Zoey_3.1.100 genome assembly. The direction of associations with p < 10^-3^ (red dotted line) is highlighted by red (hypermethylated) and blue (hypomethylated) colors. Top 15 CpGs was labeled by the neighboring genes. b) Location of top CpGs in each tissue relative to the closest transcriptional start site. The grey color in the last panel represents the location of 31,911 on the mammalian array that mapped to the genome of the Great Dane. Top CpGs were selected at p < 10^-3^. For age, top 1,000 CpGs were selected by further filtering based on z score of association with chronological age for up to 500 in a positive or negative direction. The number of selected CpGs: Age, 1,000; lifespan; 162; weight, 406. c) Boxplot of DNAm aging in CpGs located in island or outside. Labels indicate neighboring genes of the top four CpGs in each analysis. **** p<10^-4^. d) Venn diagram showing the overlap of top CpGs associated with chronological age, breed lifespan, and breed weight. The (+) and (-) signs in the table show the direction of association with each variable. e) Sector plot of EWAS of dog breed lifespan and weight. Red dotted line: p<10^-3^; blue dotted line: p>0.05; Red dots: shared CpGs; black dots: lifespan-or weight-specific changes. f) Enrichment analysis EWAS-GWAS associated genes. The heat map represents the significant results from the genomic-region based enrichment analysis between (1) the top 5% genomic regions involve in GWAS of complex traits-associated genes and (2) up to the top 1,000 hypermethylated/hypomethylated CpGs from EWAS of lifespan, adult weight, lifespan adjusted adult weight at breed level, and age at individual dog level, respectively. Cells are colored in grey if nominal P>0.05. The heat map color gradient is based on -log10 (hypergeometric P value). We list the GWAS study (x-axis) if that at least one enrichment P value (columns) is significant at a nominal significance level of 3.0×10^-3^. The y-axis lists GWAS index number (listed in **Supplementary Table 2**), trait name. The color band next to the trait encodes the GWAS category. Abbreviations: All=All ancestries, WHR= waist to hip ratio, EUR=European ancestry, FTD=frontal temporal dementia.

**Figure 4.**
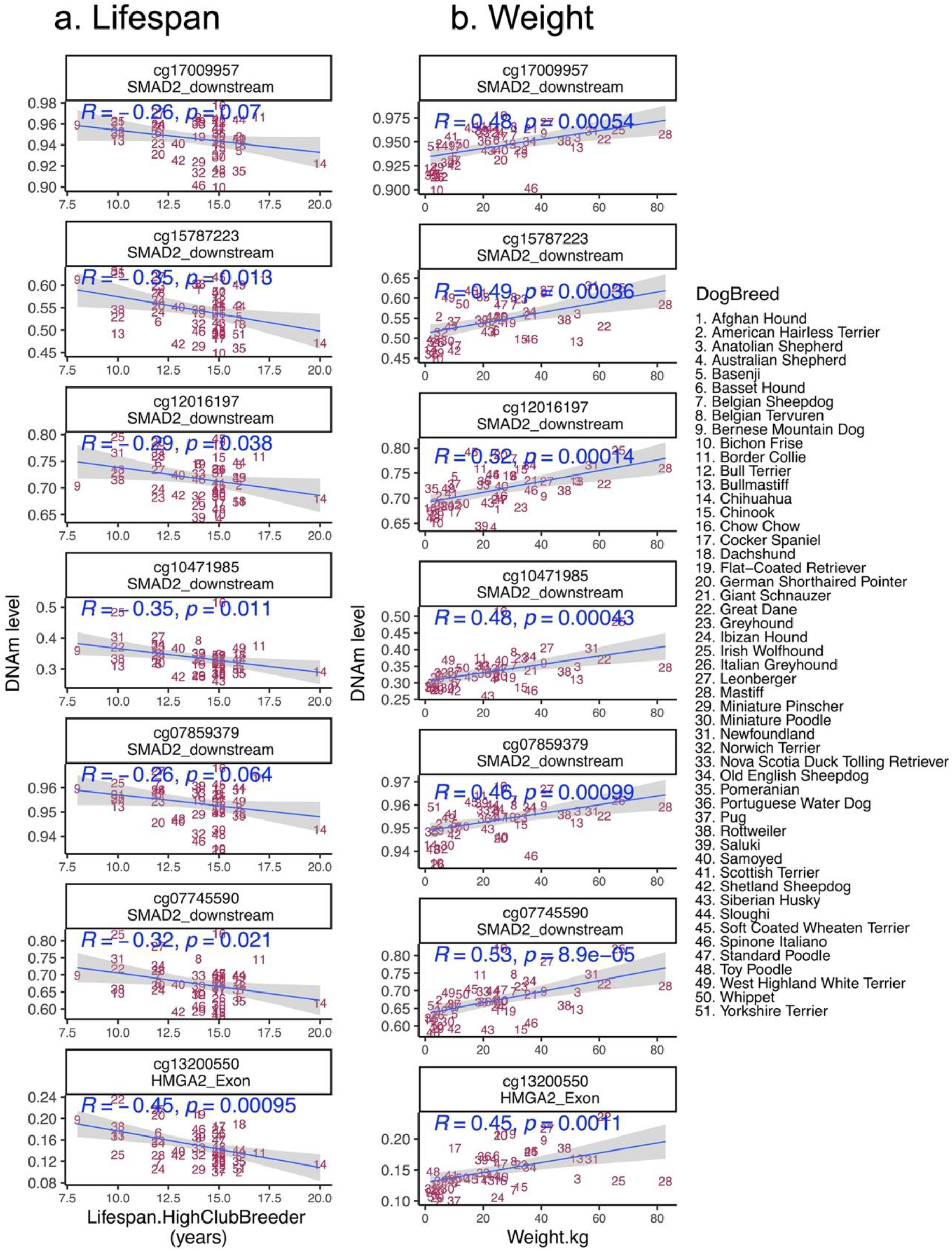
Some genes have both genetic variants and methylation patterns that relate to dog breed weight or lifespan. Scatter plots of DNAm changes with strong correlation with lifespan (a), or adult weight (b) in genes selected by GWAS of weight in dog breeds^3,16^. The title of each panel reports the CpG and an adjacent gene. The blue text inside the panel reports the Pearson correlation coefficient and the p value. The full list of CpGs that are covered on mammalian array adjacent to the top GWAS genes are reported in supplementary data.

Interestingly, age effects on cytosine methylation in dog blood correlate with those of human blood (r=0.29, **Supplementary Figure S2I**). To extend this analysis to other human tissues, we combined age effects on methylation of different human tissues via meta-analysis. We find that age effects on methylation of multiple human tissues (beyond blood) also correlate positively with that of dog blood (r=0.31, **Supplementary Figure S2A**).

Dog breeds have a unique and intriguing inverse relationship between weight and lifeexpectancy, with smaller dogs living up to two-fold longer^10^. This is contrary to observations at the interspecies level but may be due in part to domestication and artificial selection. We hypothesized that the methylation data from the 51 breeds analyzed here could reveal the identities of determinant genes and biological processes that couple growth with lifespan in domestic dogs.

Considering the moderate number of breeds at our disposal, we carried out an epigenome-wide association study (EWAS) of lifespan and weight at a nominal p-value < 10^-3^. At this level of significance, a total of 162 CpGs were associated with breed lifespan, while 406 CpGs were related to adult breed weight (**Figure 3d**). Interestingly, when CpGs with methylation changes associated with lifespan were regressed against changes with weight, a strong negative correlation (r=-0.75) was observed, consistent with the observation that bigger breeds have shorter lifespans and vice versa (**Figure 3e**). Most CpGs that have a strong positive correlation with either one of these variables, have a negative correlation with the other (**Supplementary Figure S4a**). At p-values <10^-3^, 58 CpGs were related to both lifespan and weight, but in opposing directions. Thus, this subset of CpGs may hold information regarding biological processes that underlie the intriguing inverse relationship of lifespan and weight in dog breeds. Some of the loci with this CpG methylation pattern include genes such as *KAZN* and *TBCA* which encode proteins that interact with structural and cytoskeletal proteins (desmosomes, actin and tubulin) and *NRF1,* a transcription factor that regulates expression of respiratory genes (**Supplementary Figure 4a**). An enrichment analyses of genes proximal to these 58 CpGs indicates the involvement of glycogenolysis through the *PYGM* gene (muscle-associated glycogen phosphorylase) (**Supplementary Figure S3**). Only two CpGs (one upstream of *FHL1,* the other in an intron of *MLLT3)* also appear to relate to chronological age of dogs in this study (**Figure 3d**).

We identified 34 CpGs with methylation changes that are significantly correlated with lifespan (p<10^-3^) but not with breed average weight (p>0.05) (**Supplementary Figure S4b, c**). These included CpGs in loci containing the *POLG*, *ARID5B*, *RTL1*, and *GRIA2* genes (**Supplementary Figure S4c**). Unsurprisingly, these were not significantly enriched for any pathways, as the number of genes is too low. Conversely, we identified 90 CpGs with methylation changes that correlated with breed average weight (p<10^-3^), but not lifespan (p>0.05) and observed that these CpGs are located near genes encoding transcription factors that regulate expression of genes involved in cell cycle, cell growth and adipogenesis (**Supplementary Figure S3 and S4b**). Interestingly, previously identified body size genes^15,16^, including *IGF1, LCORL*, *HMGA2*, *GHR* are not identified in the EWAS scan. Loci very specifically associated with body mass such as *IGSF1* and *ASCL4* are similarly not highlighted. That is, perhaps, not surprising as the causative variants identified so far are changes in regulatory regions and change expression levels *(ESR),* or are protein truncating *(LCORL)*^15^. This suggests that epigenetic effects are not associated with size variance for at least some classes of body size genes.

### Dog EWAS versus human GWAS

To uncover potential correlations between evolutionarily conserved cytosine variants in dogs and known human traits, the proximal genomic regions of the top positive and negative (up to 1000) CpGs identified in the EWAS described above were overlaid with the top 5% of genes that were associated with a several human traits according to Genome Wide Association Studies (GWAS, **Methods**). Overlaps were found between genes associated with body fat distribution and all the EWAS traits. For instance, there was overlap between genes implicated by a GWAS of waist-to-hip ratio in humans and those identified in our EWAS of dog breed lifespan (P=6.0×10^-8^), breed weight (P=3.0×10^-3^) and chronological age (P=3.0×10^-16^) (**Figure 3f, Supplementary Data 2**). Strikingly, genes implicated by the EWAS of dog breed lifespan overlapped with those implicated by GWAS for human longevity status (defined as longevity>85) (P=2.0×10^-3^), with overlapping genes including *WNT3* and *DACH1* and more reported in **Supplementary Data 2**. The enrichment was not confounded by breed average weight, as the overlap remained nominally significant (P=0.02) for our weight-adjusted estimate of breed lifespan (LifespanAdjWeight). The genes identified by the EWAS of lifespan also overlapped (p=0.02) with genes identified by GWAS of intrinsic epigenetic age acceleration (IEAA, derived from the human pan tissue epigenetic clock^17^). Genes implicated by the EWAS of chronological age in dogs overlapped significantly with genes implicated by GWAS of human maternal longevity (based on lifespan of mothers) (P=9.0×10^-4^ e.g. *HOXC4* and *DLX-6),* age at menarche, educational attainment, frontotemporal dementia, age-related macular degeneration, GWAS results for human DNA methylation-based estimators of granulocyte abundance^18,19^ and mortality risk (DNAmGrimAge acceleration)^12^. We note that the significance levels are not adjusted for multiple comparisons and therefore the results must be interpreted with caution.

### DNAm patterns in genes identified by GWAS of dog body weight

Previous GWAS studies highlighted five genes (*IGF1, HMGA2, SMAD2, LCORL, IGSF1*) with variants that explain around 60% of body weight variability in dogs^3,15^. The mammalian methylation array contains 34 probes adjacent to different regions of *SMAD2, IGF1, IGSF1,* and *HMGA2* (Supplementary data). Interestingly, a subset of six CpGs downstream of *SMAD2* were strongly related to adult weight, but weakly correlated with breed lifespan (Figure 5). There was also one CpG in an exon of *HMGA2* that relates to both weight and lifespan (Figure 5). The few covered mammalian probes adjacent to *IGF1* and *IGSF1* did not relate to these breed characteristics.

## DISCUSSION

Epigenetic clocks were first developed for studies of human aging traits, and in a very short time their reported use covered a large swath of medical and scientific research areas^11,20^. It quickly became clear that these clocks were able to predict chronological age with remarkable accuracy while also capturing many important features of the biological aging process. They readily found their way in biomedical applications, including the measure of epigenetic age in human clinical trials. More recently developed human epigenetic clocks are able to predict future healthspan and lifespan in a manner believed to be independent of all other established aging biomarkers^11,21^. Several mouse epigenetic clocks have since been developed and successfully validated against putative longevity treatments or genetic interventions such as rapamycin, caloric restriction and growth hormone receptor knockout models. Each of these treatments demonstrate a significant regression of epigenetic aging in mice^22–27^. These observations consolidate the notion that epigenetic clocks can be used to rapidly test the effectiveness of anti-aging interventions. Further, recent results demonstrating the presence of universal mammalian and dual-species clocks provide evidence that interventions which reverse epigenetic age in mammalian animal models may indeed translate successfully to humans, and vice versa.

There is no guarantee that an intervention which reverses epigenetic age in an animal model will successfully translate to humans, and vice versa. The likelihood of success will be increased substantially if (i) a suitable animal model for human aging is used, and (ii) a common measure of age (age equivalence) between the selected animal and humans is available, such as the domestic dog. To create a common estimator of age between mammals, however, requires that the species barrier be breached. In this study, that was achieved by the development of a mammalian DNA methylation array that profiles 36 thousand CpGs embedded within sequences that are conserved across mammals. This array platform has been used to develop epigenetic clocks for cats and many other mammalian species^14^ Most importantly, we succeeded in developing dual species clocks (human-dog clocks) that relate relative age to cytosine methylation via a log-linear transformation. These dual-species clocks can accurately estimate chronological age of dogs and humans using the same mathematical formula, highlighting yet again the common underlying mechanisms of aging between mammals. These clocks are particularly attractive as they can directly translate research findings from dogs to humans and vice versa.

With our DNA methylation data set we also developed an epigenetic estimator of average time-to-death, which is a potential estimator of mortality risk for individual dogs based on blood methylation profiles. However, this predictor requires validation. Unlike genetics, which remain unchanged through the life of an individual, DNA methylation changes with age and/or in response to environment, lifestyle and certain diseases states at a subset of loci. Hence, it would be interesting to test the performance of this estimator in prospective studies with dogs in varying states of health/illness, recovery and mortality. While our clocks were trained in blood, we expect that they lead to high age correlations in saliva or buccal samples as well. However, there could be a constant offset term (difference between DNAmAge and age) in these sample types and in other alternative sources of DNA that enable more routine and convenient sample collection.

Canine CpGs with methylation changes that correlate with breed average weight were found to be in close proximity to genes involved in adipogenesis, and age-related canine CpGs also exhibit a highly significant overlap with waist-to-hip ratio in humans. The finding that CpGs which gain methylation with canine age are located in CpG islands near targets of polycomb repressor complex 2 and developmental genes is consistent with findings in humans^28–30^. This highlights the increasingly frequent observation that the process of development is connected to epigenetic aging. There is no further empirical data to develop a more focused hypothesis, but the availability of epigenetic clocks promises to remedy this. Previous GWAS highlighted several genes with variants that explains around 60% of variance in the weight of dogs^3,16^. Interestingly, the methylation pattern in several CpGs adjacent to these genes (e.g. *SMAD2, HMGA2*) also related to weight of dog breeds. However, these CpGs had a weak correlation with lifespan, thus, probably do not underlie the inverse relationship of weight and lifespan in dog breeds.

Further similarity between epigenetics of dogs and humans is evident by the overlapping of age-related canine genes and those implicated in human maternal longevity, age at menarche, educational attainment, frontotemporal dementia, age-related macular degeneration and mortality risk as measured by GrimAge clock^12^ Genes implicated by the EWAS of dog breed lifespan overlapped with those implicated by GWAS for human longevity status (defined as longevity>85) with overlapping genes including *WNT3* and *DACH1* and more reported in Supplementary Data 2.

It is of particular interest that age-related CpGs of dogs and humans across different tissues are well-correlated; supporting the appropriateness of dogs as models for human aging. The inverse relationship between breed size and lifespan, an intriguing feature of dogs, could prove enlightening for humans if it were understood at the molecular level. The clear and substantial number of DNA methylated loci in canine DNA with opposing correlations with regards increasing weight and decreasing lifespan provides the first clue and opportunity to investigate this hitherto unexplained phenomenon.

Several articles have shown that aging effects on methylation levels are conserved at specific locations that play a role in mammalian development (reviewed in^11,31^). For example, our prior work on dogs, wolves and humans demonstrated that age associations of syntenic CpGs were conserved between canids and humans even though the data were generated on different platforms^7^ To create a robust estimator of epigenetic age for canines, humans (and many other species), we used a DNA methylation array that profiles CpGs embedded within sequences that are conserved across mammals^9^. This array platform has been used recently to develop epigenetic clocks for cats and other mammalian species^14,32^.

Recently, Wang et al (2020) developed an oligo-capture system to characterize the canine DNA methylome, targeting syntenic regions of the genome^4^. In this work, the authors present an epigenetic clock that was trained in one species (e.g. dog) but leads to a moderately high correlation in another species (e.g. dogs-to-mice, age correlation r=0.73). By contrast, our human-dog clocks were trained in both species (humans and dogs) at the same time, which may explain the substantially higher age correlation (r=0.97). Other factors such as validation schemes, sample sizes, sample preparation methods, and methylation assay technical variability may impact the results as well.

Collectively, the successful development of epigenetic clocks for dogs and the dual-species clocks, in particular, underlies the universality of the epigenetic aging process. It demonstrates that, at least at the DNA level, there are remarkable similarities in the aging process between dogs and humans. Finally, our EWAS results highlight gene regions that may underlie the inverse relationship between breed size and lifespan.

## MATERIALS AND METHODS

### Materials

In total, we analyzed DNA from n=565 blood samples (dogs) from 51 breeds. Samples were provided by researchers at the National Human Genome Research Institute (NHGRI) and collection was approved by the Animal Care and Use Committee of the Intramural Program of NHGRI at the National Institutes of Health (Protocol 8329254). The median dog age was 5.6 years (range= 0.1-17 years) (**Supplementary Table 1**). DNA isolated blood samples from dogs were used to build an epigenetic clock for dogs. To build human-dog clocks we added n=1207 human tissue samples to the training data.

### Lifespan and breed characteristics

For each phenotype, we used the average of the standard breed (male + female average). Standard breed weights (SBW), height (SBH) and life span were obtained from several sources: weights and height previously listed in^16,33^, although they were updated if weights specified by the AKC^10^ were different. If the AKC did not specify SBW, SBH or life span, we used data from Atlas of Dog Breeds of the World^34^ SBW, SBH and life span were applied to all samples from the same breed. Lifespan estimates are available for all dogs within the 51 breeds. Since Bull Terrier and Dachshund breeds have both standard and miniature sizes, the adult weights differ between these two sizes. Therefore, we assigned weight as missing for those breeds in this analysis.

### Human tissue samples

To build the human-dog clock, we analyzed previously generated methylation data from n=1207 human tissue samples (adipose, blood, bone marrow, dermis, epidermis, heart, keratinocytes, fibroblasts, kidney, liver, lung, lymph node, muscle, pituitary, skin, spleen) from individuals whose ages ranged from 0 to 93. The tissue samples came from three sources: tissue and organ samples came from the National NeuroAIDS Tissue Consortium^35^; blood samples from the Cape Town Adolescent Antiretroviral Cohort study^36^; blood, skin and other primary cells were provided by Kenneth Raj^37^. All were obtained with Institutional Review Board approval (IRB#15-001454, IRB#16-000471, IRB#18-000315, IRB#16-002028).

### DNA methylation

All data were generated on the platform (HorvathMammalMethylChip40). The mammalian array provides high coverage (over one thousand-fold) of highly conserved CpGs in mammals, but focuses only on 36k CpGs that are highly conserved across mammals. Out of 37,492 CpGs on the array, 35,988 probes were chosen to assess cytosine DNA methylation levels in mammalian species^9^. The particular subset of species for each probe is provided in the chip manifest file which can be found at Gene Expression Omnibus (GEO) at NCBI as platform GPL28271. The SeSaMe normalization method was used to define beta values for each probe^38^. Genome coordinates for different dog breeds have been posted on Github as detailed in the section on Data Availability.

### Penalized Regression models

We developed the six different epigenetic clocks for dogs by regressing chronological age on all CpGs. Age was not transformed. We used all tissues for the pan-tissue clock. We restricted the analysis to blood, liver, and brain tissue for the blood, liver, and brain tissue clocks, respectively. Penalized regression models were created with the R function “glmnet”^39^. We investigated models produced by both “elastic net” regression (alpha=0.5). The optimal penalty parameters in all cases were determined automatically by using a 10-fold internal crossvalidation (cv.glmnet) on the training set. By definition, the alpha value for the elastic net regression was set to 0.5 (midpoint between Ridge and Lasso type regression) and was not optimized for model performance. We performed a cross-validation scheme for arriving at unbiased (or at least less biased) estimates of the accuracy of the different DNAm based age estimators. One type consisted of leaving out a single sample (LOOCV) from the regression, predicting an age for that sample, and iterating over all samples.

### Relative age estimation

To introduce biological meaning into age estimates of dogs and humans with very different lifespans, as well as to overcome the inevitable skewing due to unequal distribution of data points from dogs and humans across age range, relative age estimation was made using the formula: Relative age= Age/maxLifespan where the maximum lifespan for dogs and humans were set to 24 years and 122.5 years, respectively.

### Epigenome wide association studies (EWAS) of age, lifespans and weight

EWAS was performed in each tissue separately using the R function “standardScreeningNumericTrait” from the “WGCNA” R package^40^.

### Epigenetic clock for average time to death

We did not have follow-up information (time to death) available for individual dogs in our study. To create a surrogate variable for this important endpoint, we leveraged two other variables: 1) the upper limit of lifespans estimated from the American Kennel Club and other registering bodies (variable name “Lifespan.HighClubBreeder”^10^) and 2) the chronological age at the time of the blood draw. For each dog, we defined average time to death (AverageTimeToDeath) as the difference between Lifespan.HighClubBreeder and chronological age.

To assess the accuracy of the elastic net regression models, we used leave-one-breed-out (LOBO) cross validation. The LOBO cross validation approach trained each model on all but one breeder. The “left out” breed was then used as a test set. The LOBO approach assesses how well the penalized regression models generalize to breeds that were not part of the training data. To ensure unbiased estimates of accuracy, all aspects of the model fitting (including prefiltering of the CpG) were only conducted in the training data for the LOBO analysis. We fitted the glmnet model to the top 5000 CpGs with the most significant median Z score (lifespan correlation test) in the training data. We average the methylome for each breed and performed EWAS of lifespan (N=50 breeds in training dataset) to select the top 5000 CpGs.

### Phylogenetically independence contrasts analysis

We computed phylogenetically independence contrasts (PICs) traits on lifespan, adult weight, DNAm biomarkers based on the R package “ape”. For the variable with individual level such as age-adjusted DNAmAverageTimeToDeath, we first averaged the variable at each breed then applied PICs analysis on the breed level measure.

### EWAS-GWAS based overlap analysis

Our EWAS-GWAS based overlap analysis related gene sets found by our EWAS of age with the gene sets identified by published large-scale GWAS of various phenotypes, including body fat distribution, lipid panel outcomes, metabolic outcomes, neurological diseases, six DNAm based biomarkers, and other age-related traits (**Supplementary Table 2**). Data from 97 GWAS were utilized. The six DNAm biomarkers included four epigenetic age acceleration measures derived from 1) Horvath’s pan-tissue epigenetic age adjusted for age-related blood cell counts referred to as intrinsic epigenetic age acceleration (IEAA)^17,19^, 2) Hannum’s blood-based DNAm age^41^; 3) DNAmPhenoAge^42^; and 4) the mortality risk estimator DNAmGrimAge^12^, along with DNAm-based estimates of blood cell counts and plasminogen activator inhibitor 1(PAI1) levels^12^ For each GWAS result, we used the MAGENTA software to calculate an overall GWAS *P*-value per gene, which is based on the most significant SNP association *P*-value within the gene boundary (+/− 50 kb) adjusted for gene size, number of SNPs per kb, and other potential confounders^43^. For each EWAS result, we studied the genomic regions from the top 1,000 CpGs hypermethylated and hypomethylated loci with dog relevant traits: lifespan, weight, lifespan adjusted weight, and chronological age, respectively. All of the top 1,000 CpGs were associated with the corresponding trait at P<0.05 in our EWAS, except those that negatively correlated with lifespan adjusted weight. For those, the top 633 CpGs had an EWAS of P<0.05. To assess the overlap with a test human trait, we selected the top 5 % genes for each GWAS trait and calculated one-sided hypergeometric P-values based on genomic regions as detailed in^44,45^. The number of background genomic regions in the hypergeometric test was based on the overlap between the entire genes in a GWAS and the entire dog genomic regions (~31k CpGs) in our mammalian array. We present the GWAS trait when its hypergeometric P-value reached a nominal significance level of 3.0×10^-3^ with any traits. We caution the reader that the analysis was not adjusted for multiple comparisons.

### URLs

American Kennel Club akc.org/dog-breeds/

AnAge, http://genomics.senescence.info/help.html#anage

UCSC genome browser: http://genome.ucsc.edu/index.html

Great Dane Assembly: CanFam_GreatDane.UMICH_Zoey_3.1.100

## Acknowledgements

The development of the dog tissue clocks was supported by the Paul G. Allen Frontiers Group (SH) and a grant from Open Philanthropy (SH). Human tissue sample collection was supported by NIH funding through the NIMH and NINDS Institutes by the following grants: Manhattan HIV Brain Bank (MHBB): U24MH100931; Texas NeuroAIDS Research Center (TNRC): U24MH100930; National Neurological AIDS Bank (NNAB): U24MH100929; California NeuroAIDS Tissue Network (CNTN): U24MH100928 Data Coordinating Center (DCC): U24MH100925. Human blood samples were supported by R21MH107327. The contents are solely the responsibility of the authors and do not necessarily represent the official view of the NNTC or NIH.

## Conflict of Interest Statement

SH is a founder of the non-profit Epigenetic Clock Development Foundation which plans to license several patents from his employer UC Regents. These patents list SH as inventor. Robert T. Brooke is the Executive Director of the Epigenetic Clock Development Foundation. The other authors declare no conflicts of interest.

## Data Availability Statement

The data will be made publicly available as part of the data release from the Mammalian Methylation Consortium. Genome annotations of these CpGs can be found on Github https://github.com/shorvath/MammalianMethylationConsortium

## Supplementary Figures

**Supplementary Figure S1.**
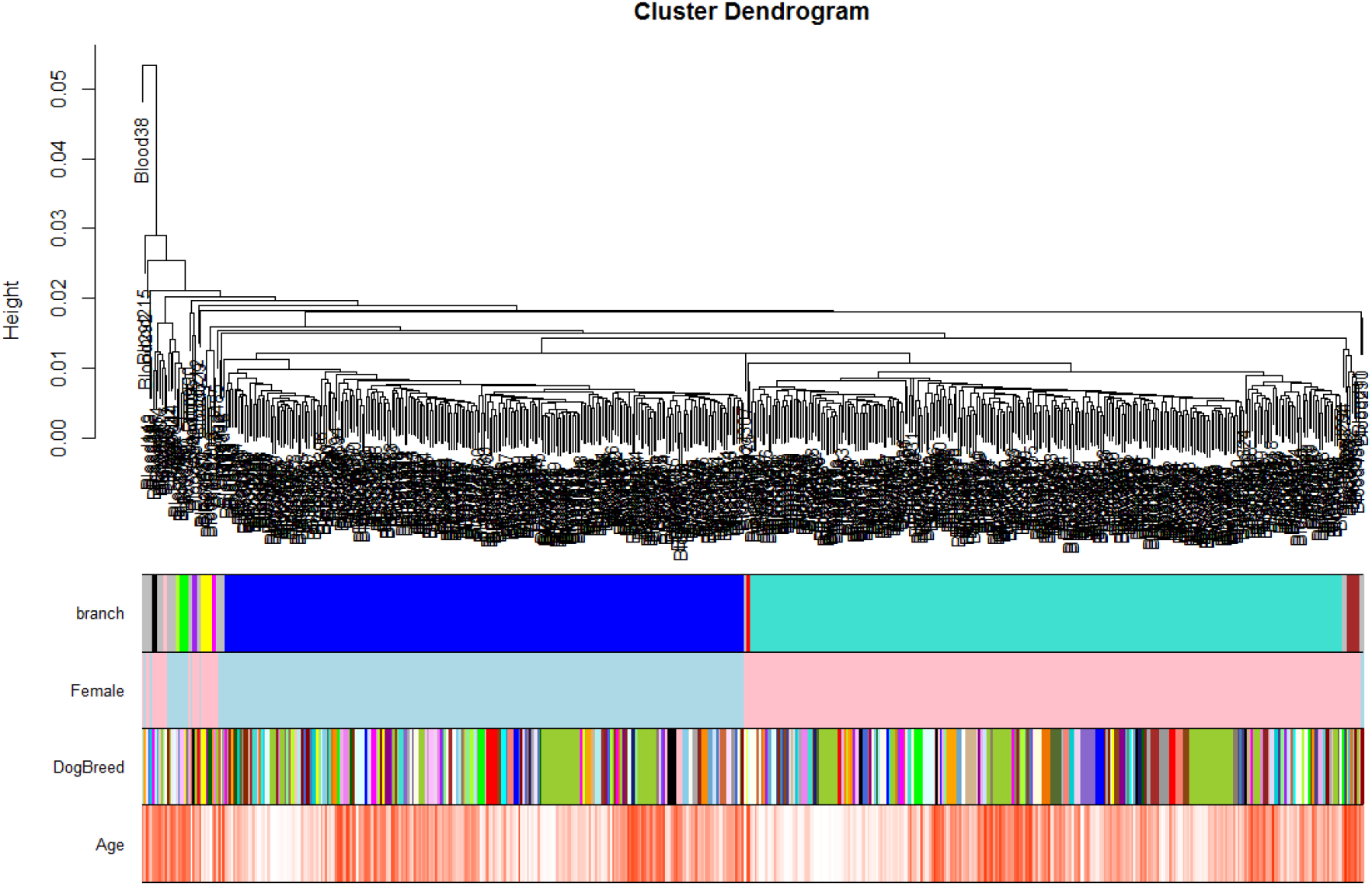
Unsupervised hierarchical clustering of blood samples from dogs. Branches largely correspond to sex as is evident from the second color band (female=pink, male=light blue). By contrast, dog breed does not lead to distinct clusters.

**Supplementary Figure S2.**
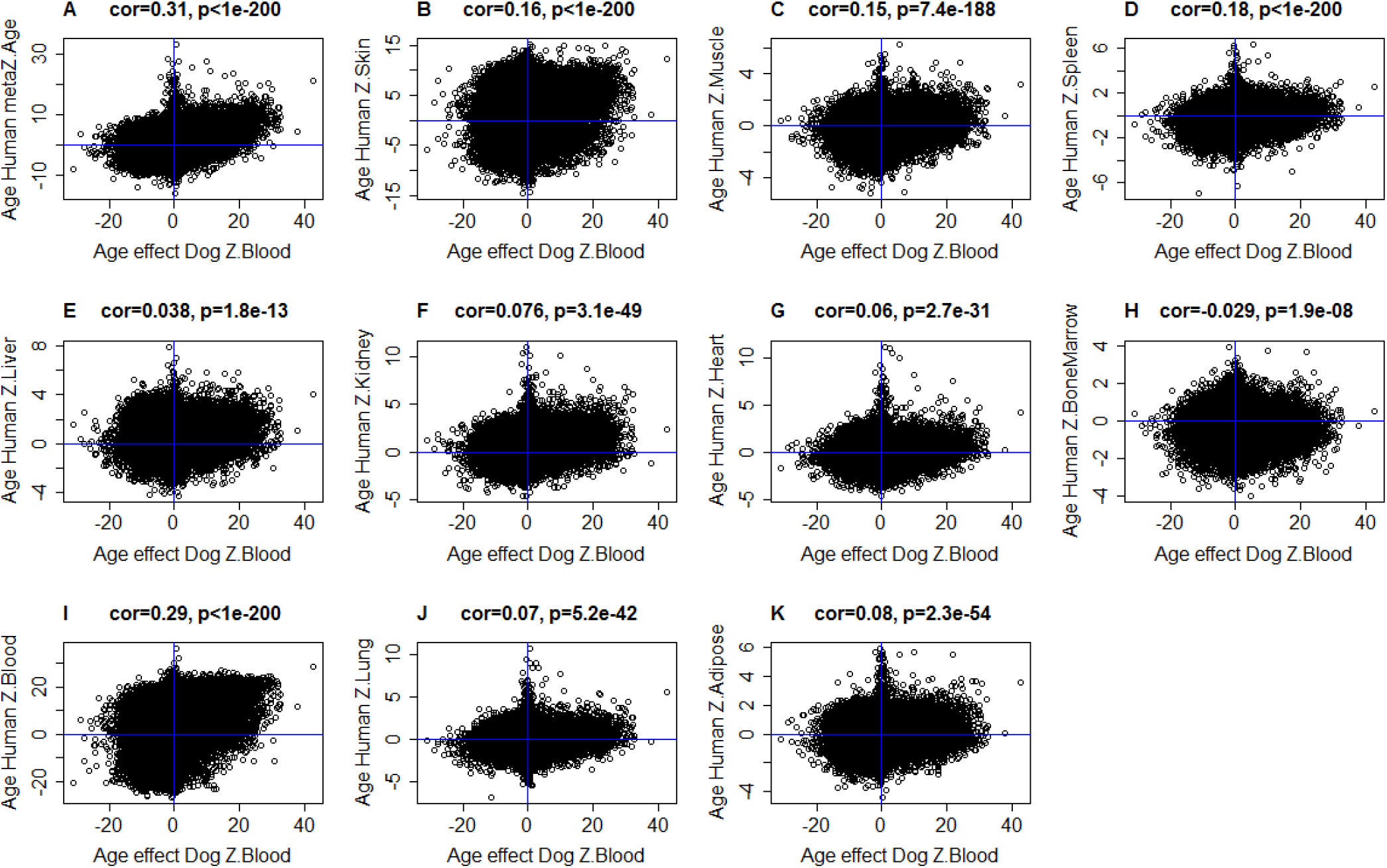
Age effects in dog blood (x-axis) versus age effects in different human tissues. a) Meta-analysis across human tissues (Stouffer’s method) was used to combine the results across human tissues. Age effects in dog blood (x-axis) versus age effects in human b) skin samples (y-axis), c) muscle, d) spleen, e) liver, f) kidney, g) heart, h) bone marrow, i) blood, j) lung, k) adipose. Each dot corresponds to a CpG on the mammalian array. Each axis reports a Z statistic from a correlation test relating a CpGs to chronological age. Positive Z statistic values correspond to a positive correlation between age and the CpG, i.e., age-related gain of methylation. The Z statistic follows a standard normal distribution under the null hypothesis (of zero correlation). A Z statistic value larger than 2 (or smaller than −2) corresponds to a (two sided) correlation test p-value less than 0.05

**Supplementary Figure S3.**
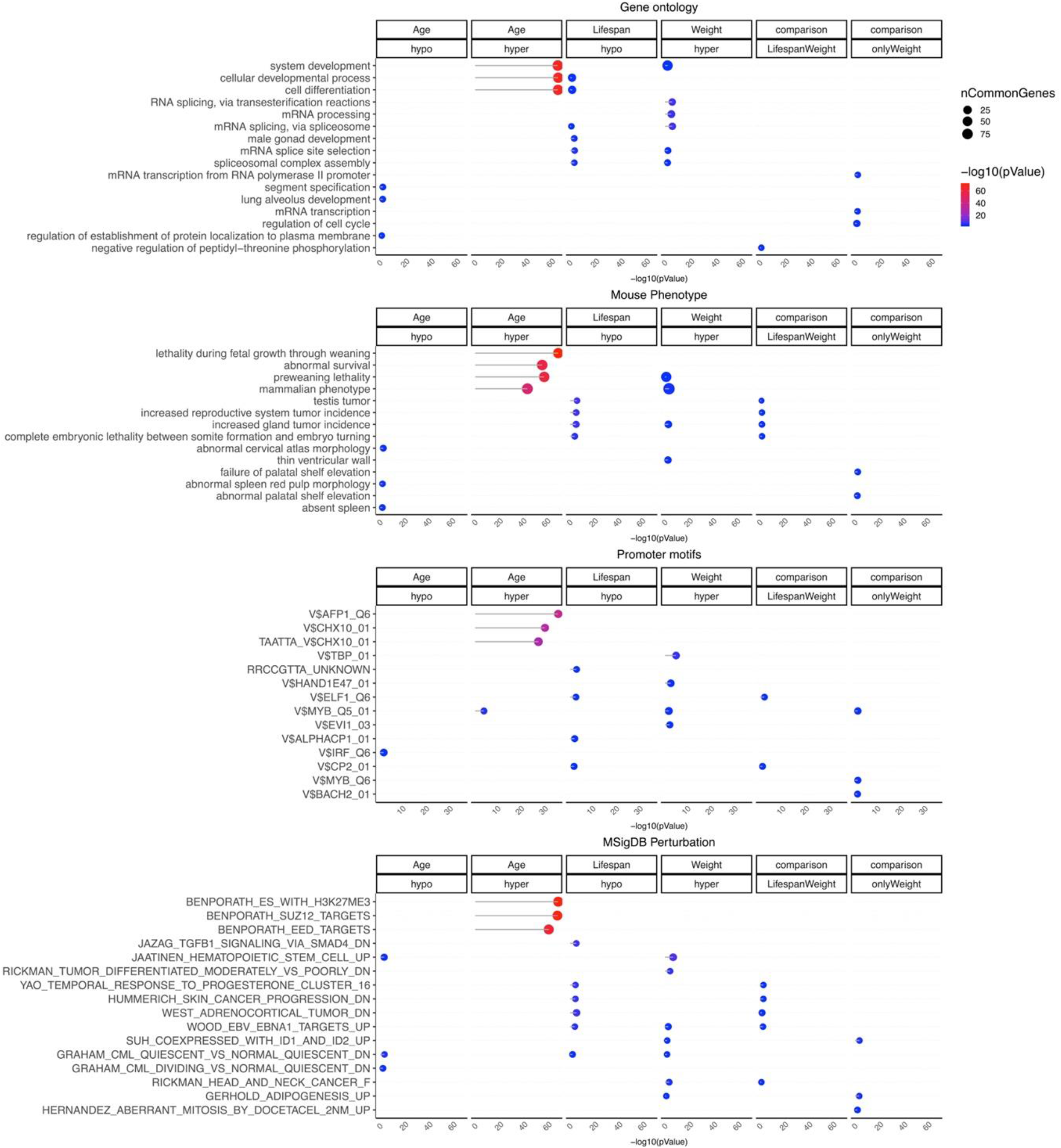
Gene set enrichment analysis of CpGs related to age, breed lifespan, and breed weight of dogs. The analysis was done using the genomic region of enrichment annotation tool^46^. The gene level enrichment was done using GREAT analysis^46^ and human Hg19 background. The background probes were limited to 20,622 probes that were mapped to the same gene in the Great Dane genome (CanFam_GreatDane.UMICH_Zoey_3.1.100 genome assembly). The figure represents enrichment at p < 10^-3^ threshold, the top two enriched datasets from each category (gene ontology, mouse phenotypes, promoter motifs, and MsigDB perturbation).

**Supplementary Figure S4.**
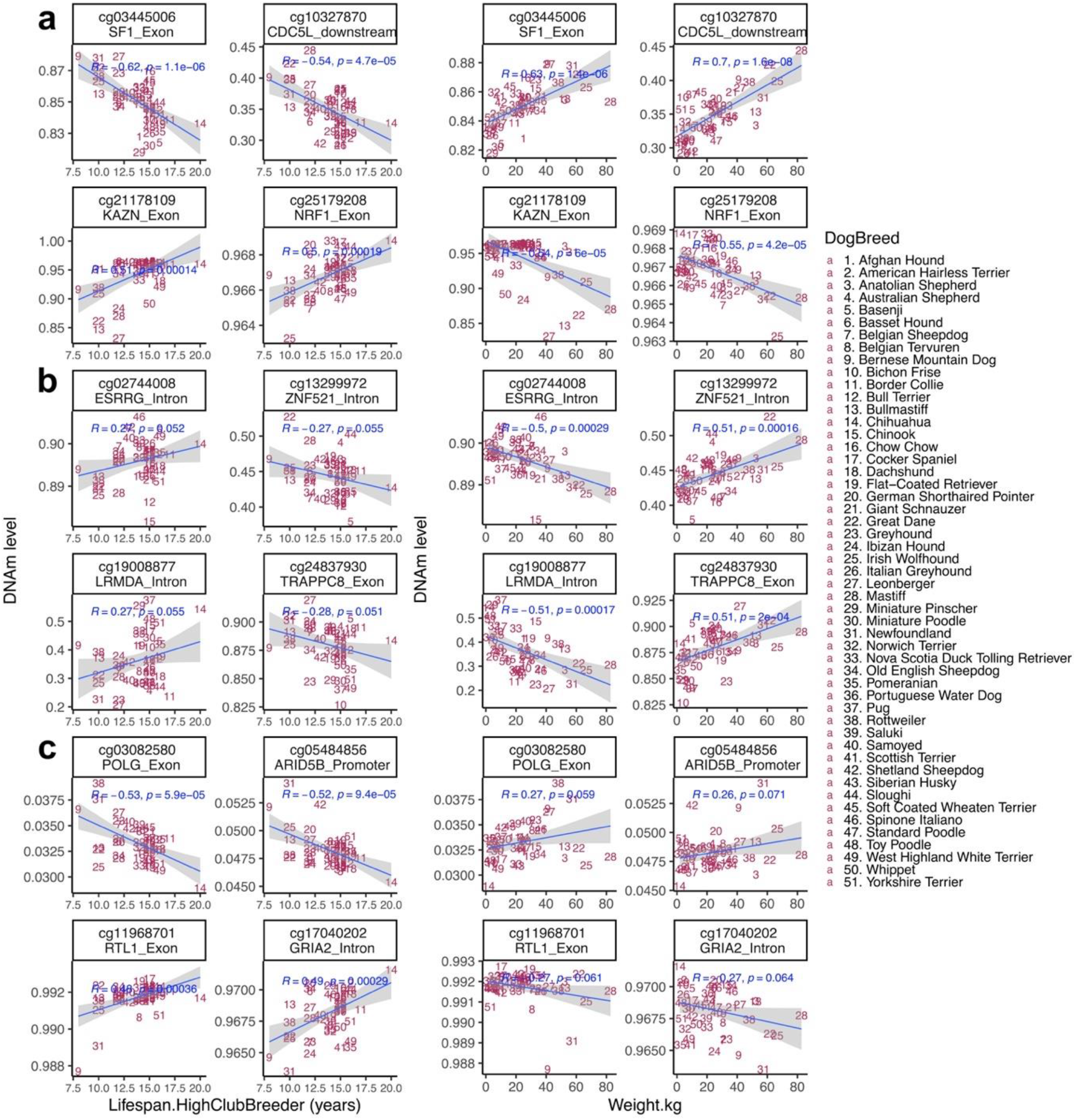
CpGs that relate to breed lifespan or average breed weight. Select CpGs that correlate significantly with a) breed weight and lifespan, b) breed weight but not lifespan, c) breed lifespan but not weight. The two left most columns depict scatter plots between breed lifespan (x-axis) and CpG methylation levels (y-axis). The two right most columns depict scatter plots between average breed weight (x-axis) and methylation levels. The title of each panel reports the CpG and an adjacent gene. The blue text inside the panel reports the Pearson correlation coefficient and the p value.

**Supplementary Table 1.** Description of the 51 pure dog breeds.

| DogBreed | Abbr. | N | No.Female | Mean Age | Min.Age | Max.Age | meanMeth | Average Lifespan | Lifespan.LowAKC | Lifespan.HighAKC | Lifespan.LowEncyclopedia | Lifespan.HighEncyclopedia | Lifespan.LowClubBreeder | Lifespan.HighClubBreeder | Weight.kg | Height.cm |
| --- | --- | --- | --- | --- | --- | --- | --- | --- | --- | --- | --- | --- | --- | --- | --- | --- |
| Afghan Hound | AfgH | 10 | 5 | 6.12 | 1.4 | 13.2 | 0.629 | 13 | 12 | 18 | 12 | 14 | 12 | 14 | 24.95 | 66.04 |
| American Hairless Terrier | AmHT | 9 | 6 | 4.28 | 0.8 | 12.6 | 0.627 | 15 | 14 | 16 | 14 | 16 | 14 | 16 | 4.76 | 35.56 |
| Anatolian Shepherd | AntS | 10 | 5 | 5.4 | 0.7 | 9 | 0.631 | 12 | 11 | 13 | 10 | 11 | 13 | 15 | 52.50 | 76.00 |
| Australian Shepherd | AstS | 9 | 4 | 6.33 | 1.3 | 13.2 | 0.631 | 13.5 | 12 | 15 | 12 | 13 | 12 | 15 | 23.50 | 49.50 |
| Basenji | Bsnj | 9 | 5 | 5.46 | 0.8 | 13.9 | 0.628 | 14 | 13 | 14 | 12 | 12 | 12 | 16 | 10.21 | 42.00 |
| Basset Hound | BssH | 10 | 5 | 5.13 | 0.5 | 10.4 | 0.630 | 13.5 | 12 | 13 | 12 | 12 | 10 | 12 | 24.50 | 33.00 |
| Belgian Sheepdog | BlgS | 10 | 6 | 7.3 | 1.1 | 13.1 | 0.623 | 12 | 12 | 14 | 13 | 14 | 10 | 12 | 30.62 | 60.96 |
| Belgian Tervuren | BlgT | 10 | 5 | 10.4 | 1.2 | 16.6 | 0.627 | 13 | 12 | 14 | 12 | 14 | 12 | 14 | 30.62 | 60.96 |
| Bernese Mountain Dog | BrMD | 11 | 6 | 6.76 | 1.3 | 14.2 | 0.629 | 7 | 7 | 10 | 10 | 12 | 6 | 8 | 41.00 | 64.45 |
| Bichon Frise | BchF | 7 | 4 | 8.26 | 2.5 | 14.4 | 0.630 | 14.5 | 14 | 15 | 14 | 14 | 12 | 15 | 4.00 | 27.00 |
| Border Collie | BrdC | 10 | 5 | 7.18 | 1.3 | 12.4 | 0.630 | 13.5 | 12 | 15 | 12 | 14 | 10 | 17 | 19.00 | 50.00 |
| Bull Terrier | BlIT | 10 | 7 | 6.15 | 1 | 13.9 | 0.626 | 12.5 | 12 | 13 | 11 | 13 | 10 | 15 | NA | NA |
| Bullmastiff | BlIm | 10 | 7 | 4.08 | 1 | 9.1 | 0.630 | 9 | 7 | 9 | 10 | 12 | 8 | 10 | 52.16 | 64.77 |
| Chihuahua | Chhh | 6 | 4 | 4.48 | 1.9 | 6.9 | 0.630 | 15 | 14 | 16 | 12 | 14 | 12 | 20 | 1.81 | 19.05 |
| Chinook | Chnk | 6 | 3 | 7.95 | 0.4 | 14.3 | 0.628 | 13.5 | 12 | 15 | 13 | 14 | 13 | 15 | 33.00 | 58.50 |
| Chow Chow | ChwC | 10 | 5 | 6.16 | 0.7 | 11 | 0.627 | 10 | 8 | 12 | 11 | 12 | 9 | 15 | 25.90 | 51.00 |
| Cocker Spaniel | CckS | 8 | 4 | 6.16 | 1.1 | 14 | 0.630 | 14 | 10 | 14 | 13 | 14 | 12 | 15 | 10.21 | 36.83 |
| Dachshund | Dchs | 10 | 3 | 6.73 | 0.8 | 13.6 | 0.630 | 15.5 | 12 | 16 | 14 | 17 | 12 | 16 | NA | NA |
| Flat-Coated Retriever | F-CR | 10 | 8 | 7.32 | 1.3 | 13 | 0.630 | 9 | 8 | 10 | 12 | 14 | 8 | 14 | 29.00 | 57.00 |
| German Shorthaired Pointer | GrSP | 9 | 4 | 5.8 | 0.8 | 13.5 | 0.630 | 11 | 10 | 12 | 12 | 14 | 10 | 12 | 26.00 | 59.50 |
| Giant Schnauzer | GntS | 10 | 4 | 7.02 | 1.4 | 14.6 | 0.631 | 13.5 | 12 | 15 | 11 | 12 | 12 | 15 | 36.50 | 65.00 |
| Great Dane | GrtD | 7 | 4 | 3.57 | 0.4 | 8.6 | 0.632 | 7 | 7 | 10 | 10 | 10 | 8 | 10 | 61.69 | 78.74 |
| Greyhound | Gryh | 10 | 6 | 5.92 | 1.6 | 10.8 | 0.630 | 11.5 | 10 | 13 | 10 | 12 | 10 | 12 | 33.00 | 72.00 |
| Ibizan Hound | IbzH | 10 | 5 | 6.35 | 0.9 | 12.2 | 0.629 | 12.5 | 11 | 14 | 12 | 12 | 10 | 12 | 25.00 | 66.00 |
| Irish Wolfhound | IrsW | 11 | 6 | 5.86 | 1.3 | 11.4 | 0.628 | 8 | 6 | 8 | 11 | 11 | 6 | 10 | 66.63 | 78.74 |
| Italian Greyhound | ItlG | 10 | 5 | 5.7 | 0.6 | 12.5 | 0.627 | 14.5 | 14 | 15 | 13 | 14 | 12 | 15 | 4.30 | 35.56 |
| Leonberger | Lnbr | 8 | 5 | 5.66 | 3.1 | 9.4 | 0.632 | 8.5 | 7 | 7 | 12 | 14 | 10 | 12 | 41.96 | 72.39 |
| Mastiff | Mstf | 10 | 4 | 5.36 | 0.7 | 12.3 | 0.631 | 11 | 6 | 10 | 10 | 12 | 10 | 12 | 82.78 | 73.03 |
| Miniature Pinscher | MntrPn | 10 | 7 | 5.69 | 0.6 | 12.8 | 0.627 | 14 | 12 | 16 | 13 | 14 | 10 | 14 | 4.25 | 27.50 |
| Miniature Poodle | MntrPd | 10 | 4 | 7.53 | 1.8 | 15.6 | 0.627 | 15 | 12 | 13 | 14 | 17 | 12 | 15 | 7.71 | 31.75 |
| Newfoundland | Nwfn | 10 | 3 | 4.84 | 0.6 | 10.3 | 0.629 | 9.5 | 9 | 10 | 11 | 11 | 8 | 10 | 57.50 | 69.50 |
| Norwich Terrier | NrwT | 10 | 5 | 7.02 | 0.5 | 15 | 0.628 | 13 | 12 | 15 | 14 | 14 | 12 | 14 | 5.44 | 25.40 |
| Nova Scotia Duck Tolling Retriever | NSDTR | 10 | 7 | 5.36 | 0.3 | 10.9 | 0.633 | 13 | 12 | 14 | 12 | 14 | 10 | 14 | 20.00 | 48.00 |
| Old English Sheepdog | OIES | 9 | 5 | 6.84 | 1.7 | 11.8 | 0.627 | 11 | 10 | 12 | 12 | 13 | 10 | 12 | 36.00 | 56.00 |
| Pomeranian | Pmrn | 10 | 5 | 3.82 | 0.5 | 13.5 | 0.628 | 14 | 12 | 16 | 15 | 15 | 12 | 16 | 2.27 | 20.00 |
| Portuguese Water Dog | PrWD | 95 | 54 | 5.46 | 0.1 | 14.5 | 0.629 | 13.5 | 11 | 13 | 12 | 14 | 12 | 15 | 20.41 | 50.80 |
| Pug | Pug | 8 | 5 | 6.35 | 0.8 | 14.3 | 0.629 | 14 | 13 | 15 | 13 | 15 | 12 | 15 | 10.00 | 27.50 |
| Rottweiler | Rttw | 10 | 2 | 6.75 | 0.7 | 13.2 | 0.631 | 9 | 9 | 10 | 11 | 12 | 8 | 10 | 48.00 | 62.50 |
| Saluki | Salk | 10 | 4 | 7.4 | 2.8 | 14.1 | 0.629 | 13 | 10 | 17 | 12 | 12 | 12 | 14 | 19.50 | 64.77 |
| Samoyed | Smyd | 9 | 7 | 6.51 | 0.4 | 14.5 | 0.632 | 13 | 12 | 14 | 12 | 12 | 12 | 13 | 26.31 | 53.98 |
| Scottish Terrier | SctT | 10 | 6 | 7.25 | 0.7 | 15.9 | 0.627 | 13.5 | 12 | 12 | 13 | 14 | 12 | 15 | 9.07 | 25.40 |
| Shetland Sheepdog | ShtS | 10 | 4 | 8.46 | 0.5 | 16.5 | 0.630 | 12.5 | 12 | 14 | 12 | 14 | 12 | 13 | 10.02 | 36.83 |
| Siberian Husky | SbrH | 10 | 7 | 4.23 | 0.3 | 9.5 | 0.630 | 13.5 | 12 | 14 | 11 | 13 | 12 | 15 | 21.55 | 54.93 |
| Sloughi | Slgh | 8 | 4 | 4.54 | 0.7 | 8.3 | 0.631 | 14 | 10 | 15 | 12 | 12 | 12 | 16 | 23.25 | 67.31 |
| Soft Coated Wheaten Terrier | SCWT | 10 | 6 | 6.97 | 0.6 | 13.7 | 0.632 | 13.5 | 12 | 14 | 13 | 14 | 12 | 15 | 15.88 | 45.72 |
| Spinone Italiano | SpnI | 9 | 3 | 4.83 | 0.7 | 10.3 | 0.629 | 11 | 10 | 12 | 12 | 13 | 12 | 14 | 36.50 | 65.00 |
| Standard Poodle | StnP | 10 | 5 | 7.3 | 0.4 | 14.3 | 0.630 | 13.5 | 10 | 18 | 11 | 15 | 12 | 15 | 26.08 | 38.10 |
| Toy Poodle | TyPd | 10 | 9 | 5.82 | 0.3 | 17 | 0.630 | 16 | 14 | 17 | 14 | 17 | 12 | 15 | 2.77 | 25.40 |
| West Highland White Terrier | WHWT | 10 | 6 | 7.81 | 1.2 | 14.5 | 0.628 | 14 | 13 | 15 | 14 | 14 | 12 | 16 | 8.00 | 26.50 |
| Whippet | Whpp | 10 | 6 | 7.46 | 0.6 | 15.9 | 0.628 | 13.5 | 12 | 15 | 13 | 14 | 12 | 15 | 12.93 | 48.50 |
| Yorkshire Terrier | YrKT | 7 | 3 | 7.36 | 1 | 13.7 | 0.629 | 13 | 11 | 15 | 14 | 14 | 13 | 16 | 3.15 | 17.7 |

